# Magnetic resonance biomarkers for timely diagnostic of radiation dose-rate effects

**DOI:** 10.1101/2023.04.28.538667

**Authors:** C. Zagrean-Tuza, M. Suditu, R. C. Popescu, M. Bacalum, D. Negut, S. Vasilca, A. Hanganu, I. Fidel, D. Serafin, O. Tesileanu, I.C. Chiricuta, A. Sadet, M.A. Voda, P.R. Vasos

## Abstract

Diagnostic of radiation effects can be obtained within hours from delivery relying on spectroscopic detection of cell metabolite concentrations. Clinical and pre-clinical studies show that radiation delivery with elevated dose-rates can achieve tumor suppression while minimizing toxicity to surrounding areas. Diagnostic biomarkers detected on short timescales are needed to orient high dose-rate radiation delivery. We have designed an ^1^H magnetic resonance approach to observe metabolite concentrations, in particular Choline, Creatine, and Lactate, in order to detect radiation dose and dose-rate effects within hours from radiation delivery. The results of our metabolic profiling method in glioblastoma cells are consistent with observations from clinical studies guided by magnetic resonance spectroscopy for radiotherapy of head tumors. At 5 Gy/min we have observed increases in lactate concentrations and decreases in [Cho]/[Cr] ratios at increasing radiation doses. An increase of the radiation dose-rate to 35 Gy/min is correlated with an increase of [Cho]/[Cr] consistent with a reduction in radiation-induced oxidative effects at high dose-rates. The observed biomarkers can be translated for radiation pulse sequences optimization.

**One Sentence Summary:** Magnetic resonance biomarkers to monitor biological effectiveness within hours after radiation delivery can be optimized for glioblastoma cells and are of potential use for the design of radiotherapy with high dose-rates.

## INTRODUCTION

Timely biomarkers for the effects of radiation-based therapy are needed to guide clinical strategy. Metabolic alterations that characterize most solid cancers [1], such as increased glucose consumption and surges in lactate, are linked to the cell energy requirements [2]. These metabolic signatures are used in functional imaging to guide cancer therapy via magnetic resonance spectroscopy (MRS) or computerized positron emission tomography (PET-CT). Improved markers dedicated to timely detection of effectiveness are especially needed when radiotherapy is involved, as the balance of benefits versus toxicity is to be evaluated early on during treatment. In radiotherapy, the principal dose-limiting constraint is the biological effects of radiation dose to organs at risk surrounding tumor target volumes [3]. The prerequisites for effective markers to guide the course of therapy are: (i) yield responses on short time scales, ideally within few hours, (ii) be non-invasive, (iii) be translatable between bench and bedside to allow continuous research input for the clinic, and (iv) enable personalized treatment, being able toreflect individual metabolism. Molecular biomarkers for imaging radiotherapy effects can be rapid and non-invasive indicators of radiobiological effects in the targeted tumor and surrounding organs [4]. Endogenous magnetic resonance biomarkers are immediately translatable between *in-vitro* and *in-vivo* settings. Magnetic Resonance experiments on cells reflect the time scale on which molecular response builds up– on the order of hours - following treatment using drugs or radiation [5]. This enables early evaluation of the efficacy and the toxicity of specific therapies. When translated to non-invasive clinical MRI, molecular biomarkers are able to yield crucial information on disease evolution in real time, thus allowing treatment optimization [6]. Metabolic markers have proven utility to diagnose difficult-to-treat tumours, such as glioblastoma or glioma of various grades [7]. MR biomarker-based approaches become increasingly used as molecular sensitivity is improved in MRI by hyperpolarisation [8]. So far an attribute almost entirely restricted to imaging techniques using radioactive tracers, molecular diagnostic emerges in MRI. In this regards, molecular MRI can be used beyond the blood-brain barrier to diagnose brain tumours [9].

### High dose-rate radiation delivery

Dose-rates receive as much consideration as the dose itself in the last years, in the quest to minimize toxicity during radiotherapy. Radiotherapy with high dose-rate delivery, up to the ‘FLASH’ regime [10], reached the phase of clinical studies four years ago in a study seeking to minimize side-effects in the treatment of skin tumours. Further clinical studies extending applications to proton beams are ongoing [11]. This stage was reached based on accumulating evidence from pre-clinical studies showing that fast-paced delivery of radiation minimizes toxicity to surrounding areas while achieving equivalent tumor suppression results to that of conventional dose-rate studies [12]. However, the full molecular mechanistic basis of FLASH toxicity-sparing effects is yet to be understood. It is emerging that the FLASH effect relies not only on a threshold of the dose-rate but also of the dose, thresholds that are cell-type-specific; differences in the levels of generated reactive oxygen species (ROS) between high dose-rate delivery and conventional dose-delivery were detected in studies on mice brains [13].

Time-compressed focused laser radiation [14, 31] with powers of up to 10 PW accelerates photons and particles to the extreme limit for pulsed high-dose rate radiation delivery. The actual dose-rate for delivery depends on: (i) the energy concentrated in the duration of each pulse, and (ii) the frequency of pulse repetition. Experiments show that free-radical-induced oxidative stress differs between dose-delivery regimes pertaining to laser-accelerated protons and standard delivery [15].

A major challenge in the field of pulsed high dose-rate radiation is the long time required for structural diagnostic of the effects of toxicity and efficiency: changes in tumor size occur on the order of several weeks / months. Moreover, such changes in size do not always correlate with therapeutic effects, as effective tumor cell death can occur even months later [16]. To optimally guide the course of therapy, high dose-rate radiation needs fast diagnostic. A rapid evaluation of radiation effects - on the time scale of tens of hours - can be achieved relying on molecular detection. Radiation delivery in high dose-rate pulses requires diagnostic methods on adapted timescales to optimize protocols and to integrate high dose-rate radiotherapy schemes with chemotherapy and immunotherapy protocols. The present paper proposes magnetic resonance biomarkers to monitor the toxicity and the efficacy of radiotherapy in vivo within hours.

## RESULTS

### Development

Metabolite detection by magnetic resonance can be performed either in intact cells using high-sensitivity detection in high-field spectrometers [17], often coupled with hyperpolarization [18], or using cell lysates. We have tailored a protocol (Figure 1A) for cell growth, irradiation, lysis and metabolite extraction procedures in order to enable detection using spectrometers with intermediate sensitivity, e.g., at proton Larmor frequencies of 500 MHz (magnetic field *B*_0_ = 11.7 T). We have compared the concentrations of metabolites from cells irradiated at different doses and non-irradiated. The cells were frozen at a time point of ca 2 h following irradiation, in order to allow development of the enzyme response on time scales related to free-radical scavenging but short with respect to cell multiplication. Sufficient resolution and sensitivity to assign individual metabolites could be obtained from cultures of 5-10’000’000 cells for each spectrum (Figure 1, Supplementary materials Figure S1, S2).

**Figure 1.**
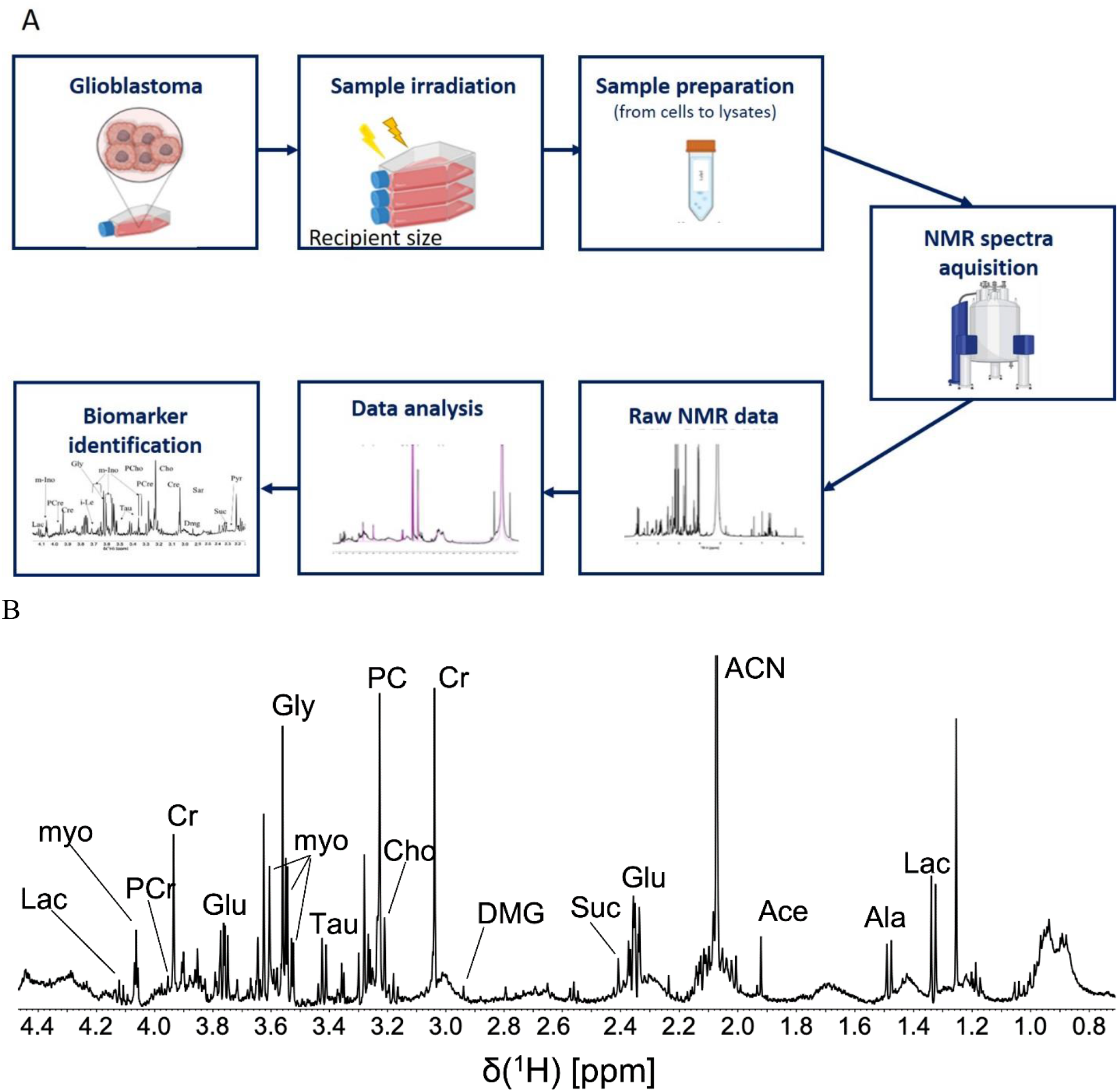
A. Outline of the metabolite extraction and observation protocol following sample irradiation involving: cell culture, sample irradiation, samples preparation and spectroscopic analysis; B. Detected NMR biomarkers using a 11.4 T spectrometer (proton Larmor frequency of 500 MHz); the displayed spectrum corresponds to non-irradiated cells. The frequency (ppm) axis discriminates chemical groups in metabolites and spectral intensities are proportional to concentrations. The assignments correspond to: Lac (lactate), Ala (alanine), Ac (acetate), Pro (proline), Suc (succinate), Dmg (dimethylglycine), Cr (creatine), PCr (phosphocreatine), Cho (choline), PCho (phosphocholine), myo (myo-inositol), Tau (taurine), Gly (glycine), Ile (isoleucine), AMP (adenosine monophosphate).

Irradiation was performed with Co-60 gamma photons from a GC-5000 research irradiator (IFIN-HH / IRASM Department) at dose-rates of 9 and 35 Gy/min, and, respectively, with X-rays at Amethyst Radiotherapy, within a water phantom placed in the 6 MV clinical irradiator, delivering radiation with a dose-rate of 5 Gy/min (equipped with a flattening filter). Given that FLASH-type effects are sought at the higher dose-rates, the dose profiles of metabolite concentration variations were pursued up to high dose values. While the cells enzymatic activity is relevantly observed in these conditions, the feasibility in terms of dose/shot in vivo obviously depends on the protective effect for normal tissue. Our optimized protocol yields metabolite signals showed in Figure 1B, where magnetic resonance proton frequencies (in ppm) allow metabolite identification and intensities allow quantifying their concentrations. The following metabolites were identified: Lac (lactate), Ala (alanine), Ac (acetate), Pro (proline), Suc (succinate), Dmg (dimethylglycine), Cr (creatine), PCr (phosphocreatine), Cho (choline), PCho (phosphocholine), myo (myo-inositol), Tau (taurine), Gly (glycine), ILE (isoleucine). The comparison between the NMR spectra from irradiated and non-irradiated cells reveals that certain important metabolite concentration ratios, reflecting anaerobic respiration and energy storage pathways, e.g., [Lac]/[Ala] and [Cho]/[Cr], change upon irradiation. Only ratios of concentrations are quantified since the number of cells, and therefore absolute concentrations of individual metabolites, differ slightly between the samples.

The comparison between the spectrum from non-irradiated cells (assigned) and the one from cells irradiated with 10 Gy photons (superimposed on top) reveals that several important metabolite ratios reflecting anaerobic respiration and energy storage pathways, e.g., [Lac]/[Ala] and [Cho]/[Cr], change upon irradiation. Only ratios of concentrations are quantified since the number of cells, and therefore absolute concentrations of individual metabolites, differ slightly between the samples.

Amongst the detected endogenous molecules, we have sought predictors for radiation effects. In the case of brain tumours, a recent review of clinical MRI studies [19] identifies a consensus marker of radiotherapy effects, which is the ratio between the concentrations of Choline and Creatine, [Cho]/[Cr]. The [Cho]/[Cr] marker can indicate within short times after radiotherapy whether treatment has been effective or if the tumor is likely to relapse within five years. Besides therapy and post-therapy assessment, it was also used for tumor classification and predictor of survival [20,21]. Choline and its derivates, phosphocholine (PC) and glycerolphosphocholine (GPC) represent markers for membrane integrity and turnover [22] and modified metabolism was observed in various cancer types, including gliomas [23]. In the context of *in-vivo* NMR [23], the value of [Cho]/[Cr] ratio may comprise of both the metabolism effect and the cellular density. Ex-vivo studies on excised tumors on different glioma types [22] and studies on different cell lines [24, 25] reported on the choline metabolism as response to radiation. In our case, [Cho]/[Cr] ratios were measured in cell samples irradiated with increasing doses using both gamma radiation from a Co-60 source in laboratory settings and X-rays delivered by a radiotherapy accelerator in the clinic. The doses employed were elevated in order to increase the likelihood of inducing FLASH-type effects. The cellular viability in terms of metabolic activity was evaluated for the cells irradiated with the high-intensity gamma source (Supplementary materials, Figure S3). Cell metabolic viability measured at the same time when samples were frozen for spectroscopy decreases monotonously with increasing doses from 50% at 10 Gy to around 35% for cells irradiated at 30 Gy.

Our results show a consistent decrease of [Cho]/[Cr] with increased radiation dose (Figure 2B) and an increase in the ratio when the radiation dose-rate is increased by a factor 4. The dependency on radiation doses was reproduced using both gamma photons from the intense Co-60 source and X-rays in the phantom placed in a clinical irradiator. The total choline (Cho, PCho, GPC) to total creatine (Cr, PCe) ratio, [tCho]/[tCr], was observed to be fairly constant with the dose for irradiated samples.

**Figure 2.**
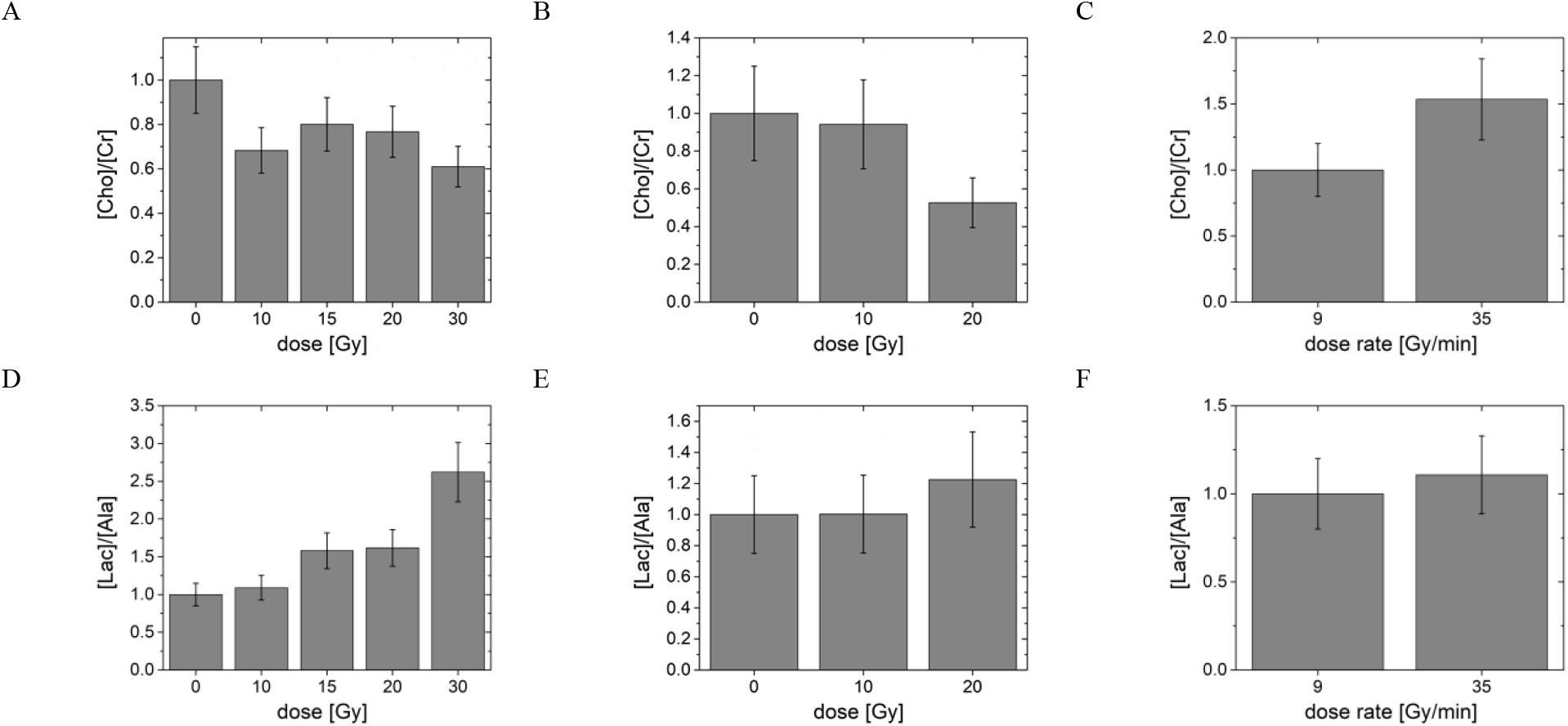
Effects of radiation on glioblastoma cells as detected by [Cho]/[Cr] biomarker (A-C) and by the [Lac]/[Ala] biomarker (D-F) within two hours after irradiation. Individual integrated signals, normalized here with respect to the inital points, are given in the Supporting Information. Dose variation effects were measured using a Co-60 intense gamma irradiator (A, D) with a dose-rate of 35 Gy/min and, respectively a water phantom setting within an X-ray clinical irradiator with a dose rate of 5 Gy/min (B, E).Dose-rate effects (C, F) were measured at a Co-60 intense gamma irradiator for a dose D = 20 Gy.

Other molecular markers such as concentrations of Lactate [Lac], Pyruvate [Pyr], etc. are effective in predicting treatment course *in-vivo* by revealing the differences in pathways related to cellular energetics between treated and non-treated tumor cells.Tumors exhibit anaerobic metabolism that is reflected in elevated levels of lactate. Normalized [Lac]concentrations were already found to depend in-vivo on the post-irradiation re-oxygenation of the tumor [26], and can be used to assess cell apoptosis after irradiation [27].

## DISCUSSION

This study presents NMR-observable metabolite ratios, namely [Cho]/[Cr] and [Lac]/[Ala], that allow optimizing and calibrating radiation therapy in cell cultures. These biomarkers are foremost of empirical value, as the metabolic pathways altered by radiation are too complex to be entirely covered by this procedure, which does not image the complete metabolome (vide infra). However, the protocol has the merits of being straightforward and based exclusively on endogenous molecules [28, 29].

Foremost, we discuss the limitations that apply to this study. The proposed metabolite detection method is demonstrated herein at intermediate field strengths; this renders it widely applicable, as a majority of laboratories have access to NMR with proton Larmor frequencies in the 400 MHz - 500 MHz range, but the ensuing limited sensitivity requires cell lysis, as in-cell signals are broadened by the viscosity of the medium and by the internal microstructure, yielding signal-to-noise ratios below what is needed for quantitative measures. Since cells are lysed after each irradiation delivery, which takes place in the culture flask where adherent cells are present in a monolayer, the signal intensities of individual metabolites are proportional to the number of cells used in each experiment. Therefore, ratios of metabolite signals or normalization to the spectral sum within the same sample rather than individual concentrations are considered. Another limitation is that the extraction protocol applied above does not transfer to the final sample the complete cells’ metabolome, due to differences in solubility between the metabolites in the chosen solvent. However, the alternate approach, direct in-cell spectroscopy, also leaves out signals of part of the metabolome: the signals that are not detected pertain to molecules featuring strong or non-specific interactions with the cell membrane or large proteins, leading to signal broadening. There are, in either case, metabolite signals missing from the spectra.

Two types of radiation effects are likely to prevail in the observed variations of metabolite ratios, induced on different timescales (Figure 3) and observed at our chosen time point for lysis and detection, within two hours after irradiation: (i) oxidative stress delivered through the creation of free radicals and ensuing fast reactions on the microsecond-to-ms range; free-radical formation triggers reactive oxygen (ROS) species and ROS effects are translated in the cell metabolome within a time lapse of minutes to hours; (ii) oxygen depletion. The later effect is related to the first as oxidative stress drives cells towards anaerobic respiration [30]. Eventually, oxygen repletion may interfere during post-irradiation sample manipulation.

**Figure 3.**
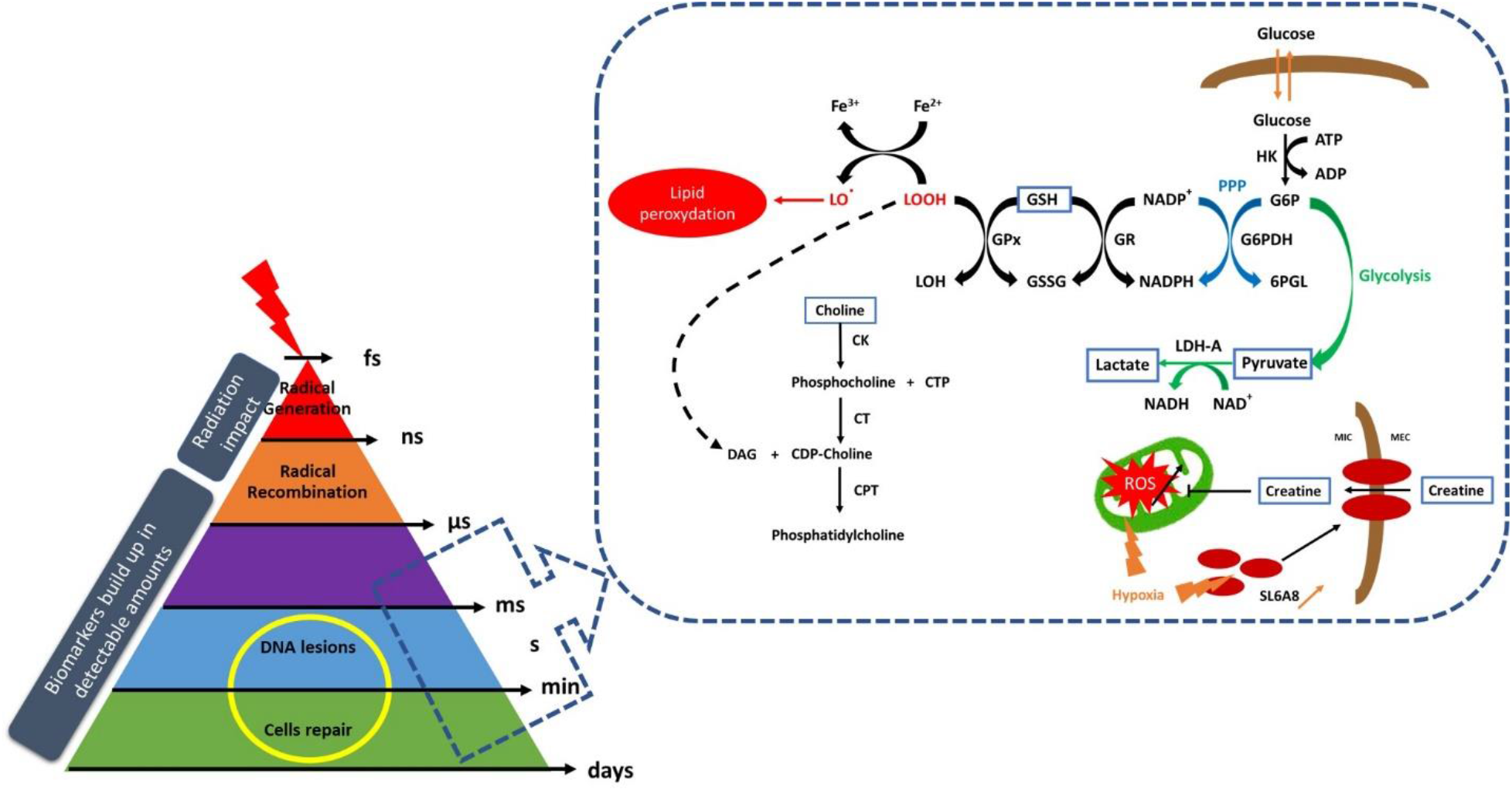
The initial radiation impact defines the time-scale beyond which cell radiation response is triggered. The response is reflected in the cellular metabolites [31], amongst which several biomarkers detected in this study are outlined.

The consistent decrease observed in [Cho/[Cr] ratios at increasing radiation doses with our metabolic profiling method in glioblastoma cells is in agreement with the behavior reviewed for clinical studies guided by MRI [32]. This marker, also known as CCrI (the choline-to-creatine index [33], is therefore applicable for observations of dose-rate effects in an empirical manner. Though the analysis in terms of metabolic pathways remains speculative given the metabolic resolution of our study, we can infer that choline depletion occur at increasing dose rates, as it was observed from the variation of [Cho] normalized to the metabolite spectral sum (Figure S4 in the Supplementary materials). Choline is a membrane constituent as well as an intermediate in the cell glycine production. Radiation-generated ROS leading to oxidative stress can induce cell death by different mechanisms, mainly apoptosis, necrosis, or, as recently underlined in the case of radiation effects, ferroptosis [34]. Antioxidants interfere with this course of events, especially since glutathione (GSH) is known to be present in large amounts in cells undergoing division, including tumor cells. Cancer cell proliferation leads to intracellular accumulation of reducing equivalents such as GSH and NADPH to control reactive oxygen species (ROS) and maintain redox balance [35]. Vitamin C is another essential micronutrient that can play the role of antioxidant and inhibit the formation of ROS. Ferroptosis mediated by oxidative processes [36], involving lipid peroxidation. In ferroptosis, choline is converted in polyunsaturated fatty acids (PUFA). Free PUFA react with iron Fe^2+^ (via a Fenton reaction) to produce phospholipid hydroperoxides, a form of lipid-based ROS [36]. Our individual metabolite signals normalized to the integrals sum indicate depletion of choline upon dose escalation.. Other reported *in-vitro* investigations of metabolite ratios for glioma cells after irradiation [24, 25] indicate an increase of [Cho]/[Cr] for irradiation with X-rays using doses of up to 16 Gy. The explanation for this trend was the enhancement of cell metabolism in response to radiation damage, damages to membrane integrity and choline leakage.

Based on the variation in the levels of other observed biomarkers such as dimethyl-glycine, assessed by NMR as well as cystine and sarcosine by Reversed Phase Liquid Chromatograpy measurements (Supplementary materials, Figure S5), it appears the metabolic pathway that leads to glycine production using choline as an intermediate may also be hindered as radiation doses are escalated. The production of γ-glutamyl-cysteine-glycine tripeptide (GSH) needed to respond to oxidative stress relies on glycine levels. As far as lactate is concerned, this is the endpoint of anaerobic respiration, accumulating as via the Warburg effect [2]. In our study, alanine (Ala) was chosen as a reference for Lac accumulation. Enhanced anaerobic respiration is a well-known effect of radiation [32], therefore the increases in [Lac]/[Ala] ratios at increased dose-rates are to be expected.

### Dose-rate effects

Using high-intensity Co-60 facility [36], we could attain a factor 4 increase in the dose-rate, approaching the regime where dose-rate effects are expected for the metabolism [37]. If, as concluded by [13], free-radical and ensuing ROS formation is lower at high dose-rates compared to conventional ones, the effects on cellular energy metabolism as reflected by [Cho]/[Cr] ratios at high dose-rates are expected to be smaller than at standard dose-rates. Our first observations (Figure 2) show this expected effect, albeit in this study the dose-rate variation was small. Anaerobic respiration is a pathway involving several oxidation steps and a final reduction of pyruvate into lactate. The oxidoreduction potential of the cells intervenes in the [Pyr]/[Lac] equilibrium via nicotinamide adenine dinucleotide level ratios [NAD+]/[NADH] [38]. This latter ratio is tuned mainly by equilibria of the most abundant antioxidant in cells, which in glioblastoma is most likely glutamate in its reduced/oxidated forms, GSH/GSSG. Herein, the [Lac]/[Ala] ratios remain relatively stable as the radiation dose-rate is increased. This either indicates that sensitivity in the dose-dependent variation of this biomarker is insufficient or that, within the competition between the anaerobic respiration effect and oxidative stress diminishing the overall reduction potential of the cells via GSG/GSSH and NADP+/NADPH intermediates [39], the change in radiation-delivery time scales may lead to small overall changes.

### Translation to ex-vivo probes and in-vivo diagnostic

The observed markers can be extended to *ex-vivo* probes. This is especially advantageous for pulsed high dose-rate radiation delivery, allowing to optimize a sequence of doses. Most of the detected molecules occur in very low concentrations (μM range) by the standards of MR detection, while in-vivo imaging most often relies on the detection of signals from abundant water protons (concentrations in the M range). Sensitivity enhancement is therefore needed for MR to detect most relevant biomarkers in radiotherapy. NMR at enhanced sensitivity using either very high-field magnets, dissolution-based Dynamic Nuclear Polarisation with magnetisation preservation, or both, affords detection of metabolites within intact cells [7, 39, 40].

Assessing treatment efficacy is often difficult in the first days or even weeks of therapy, even by using expensive imaging equipment. Non-invasive probes to assess early treatment response are of the uttermost importance for current development of healthcare. Sensitivity enhancements by factors over 10’000 for metabolite observations can be obtained by hyperpolarised Magnetic Resonance (DNP-MR) and polarisation sustaining for amino-acids and small peptides [7]. Diagnostic methods based on hyperpolarised DNP-MR for solid tumors can be used to follow in a non-invasive manner the biological effects of radiation via functional imaging [39]. These afford the detection of biomarkers triggered by radiation in vivo and in cells. Methods for detecting biomolecular transformations related to ROS formation and scavenging in real time [41] can follow cellular responses in as far as the mechanistic effects are concerned. Recent improvements in hyperpolarisation methods for DNP-MR use partly-deuterated water – HDO - to sustain magnetisation [42] followed by magnetisation transfer to biomarkers. These methods can allow observing biomarkers within intact cells, as there is little hindrance for hyperpolarised water to pass the cell membrane. The above developments reached the point where biomarkers can be observed in cell, without recurring to lysis. Thus, MR biomarkers can predict treatment outcome very early on - in a matter of tens of hours or days-from the delivery of treatment, allowing to adjust the course of radiotherapy, immunotherapy, chemotherapy or combined therapy in real time.

Dose-rates as high as Gy/ns can be obtained using particle or photon acceleration by 10 PW-class lasers [43]. Molecular markers of FLASH radiation effectiveness and toxicity can be especially useful in the case of radiobiology experiments on very short time-scales, with pulses as fast as ns, that can be achieved using high-intensity lasers [44].

## MATERIALS AND METHODS

### Cell culture and seeding prior to irradiation treatment

U-251 MG human brain glioblastoma cells (CLS Cell Line Service, Eppelheim, Germany) were cultured in Dulbecco’s Modified Eagle Medium (DMEM) supplemented with 10% fetal bovine serum and 0.1% Penicillin-Streptomycin (Biowest, Nuaillé, France). Cells were seeded in 25 mL flasks (TPP Techno Plastic Products AG, Trasadingen, Switzerland) at an amount of approximately 2’400’000 cells/flask and incubated in standard conditions of temperature and humidity during 24 h before irradiation to allow their attachment.

### Gamma irradiation

Cell culture irradiations were performed at a temperature of 25 °C, using a ^60^Co research irradiator (GC-5000, BRIT) from IRASM irradiator facility, IFIN-HH Magurele, Romania, at constant dose rates of 9 and 35 Gy/min and dose values of 10, 15, 20 and 30 Gy. Absorbed doses and dose rates were evaluated by an ethanol-chlorobenzene dosimetry system by an oscillometric readout method and indicated as absorbed doses in water.

### X-ray irradiation

A clinical irradiator at Amethyst Radiotherapy Center Otopeni, Romania, delivering X-rays (6 MV accelerating potential) at doses of 10 and 20 Gy and a dose rate of 5 Gy/min. The experimental set-up is schematically presented in Figure S6 in Supplementary materials.

### Cells processing following irradiation treatment

Following irradiation, glioblastoma cells were washed three times with ice cold PBS (phosphate buffer saline) during few minutes and detached using 1% Trypsin in PBS. Cell viability was estimated by counting method at ca 2 h after irradiation, at the same time when cell samples were stored deep-frozen for upcoming lysis and metabolite extraction procedure. On the day of the NMR experiment, the cell pellets were brought to ambient temperature and suspended in an ice-cold mixture of acetonitrile and deionized water at 1:1 volume ratio, and lysed using an ultrasound probe. The lysates were then centrifuged at 4 °C. The supernatant was carefully collected and the solvent was evaporated using a rotaevaporator. For the NMR experiments, the dry residue was dissolved in D_2_O containing 1M phosphate buffer at pH = 7.0.

### NMR measurements

NMR experiments were conducted on a Bruker Avance III spectrometer, operating at proton Larmor frequency of 500 MHz equipped with a 5 mm TX NMR rf probe. Proton metabolic profile spectrum was recorded using water suppression by saturation with excitation sculpting with gradients, the so-called zgesgppe from Bruker pulse programs library [45, 46]. A repetition time of 15 seconds was sufficient for full recovery of longitudinal magnetization between scans and a number of 1024 accumulations was set, resulting in a total measurement time of 4.75 hours. Sample temperature was maintained at 20 °C during measurements. The NMR spectra were processed and analyzed using MNova NMR (Mestrelab Research S.L.).

## Supplementary Materials

Figure S1. (A) 1H NMR spectra of the metabolite extract from the lysed GBM cells and (B) of the whole, intact cells sample.

Figure S2. (A) ^1^H NMR spectrum of the metabolite extract from a lysed sample consisting of a very large number of cells (>40 M) where glutathione (GSH) was detected and (B) Spectra detail to indicate the level of spectral resolution; a comparison between non-irradiated cells and 10 Gy-irradiated cells is provided.

Figure S3. Cell viability as a function of the dose, measured by the metabolic activity quantification (MTT test).

Figure S4. [Cho] and [Lac] as a function of radiation dose, normalized to the sum of signal integrals of the detected metabolites in order to verify the robustness of the normalization.

Figure S5. Effects of the irradiation dose (A, C) and dose-rate (B, D) on the glioblastoma cells, assessed within two hours after irradiation by HPLC: sarcosine (A, B) and cystine (C, D) are detected biomarkers. The concentrations of each analyte were normalized to the first point (unirradiated). Chromatograms were obtained using a Reversed Phase Liquid Chromatography system with o-phthaldehyde and 9-fluorenylmethoxycarbonyl chloride for online pre-column derivatization and UV-Vis detection at 241 and 330 nm.

Figure S6. Scheme of the experimental set-up for irradiation of cell samples in the clinical X-ray irradiator. The distance between the X-ray source and the flasks (each 0.1×0.07×0.04 m) was d = 1.2 m and a water phantom cube with the side of 0.4 m was used for flask fixation.

## Acknowledgments

We thank Dr Marie-Christine Vozenin, Prof. Joao Seco, and Dr Pascal Froidevaux for useful discussions.

## Funding

Romanian Ministry of Research, UEFISCDI PN-III-P4-ID-PCE-2020-2642 Romanian Ministry of Research, UEFISCDI PN-III-P4-PED-565.

Extreme Light Infrastructure Nuclear Physics (ELI-NP) Phase II, a project co-financed by the Romanian Government and the European Union through the European Regional

Development Fund and the Competitiveness Operational Programme (1/07.07.2016, ID 1334)

Romanian Ministry of Research, Equipments of National Interest (IOSIN)

Norway–Romania Grants: RO-NO-2019-0510, CTR 41/2020

## Author contributions

Conceptualization: ICC, PRV

Methodology: CZT, MS, RCP, MB, DN, SV, AS, MAV, PRV

Investigation: CZT, MS, RCP, MB, DN, SV, AH, IF, DS, AS, MAV, PRV

Visualization: CZT, SV, IF, DS, AS, MAV

Funding acquisition: OT, ICC, PRV

Project administration: OT, MS, ICC, MAV, PRV

Supervision: OT, MS, ICC, PRV

Writing – original draft: CZT, MB, SV, MAV, AS, ICC, PRV

Writing – review & editing: CZT, MB, SV, MAV, AS, ICC, PRV

