## Supplementary materials for "Magnetic resonance biomarkers for timely diagnostic of radiation dose-rate effects"

(A)

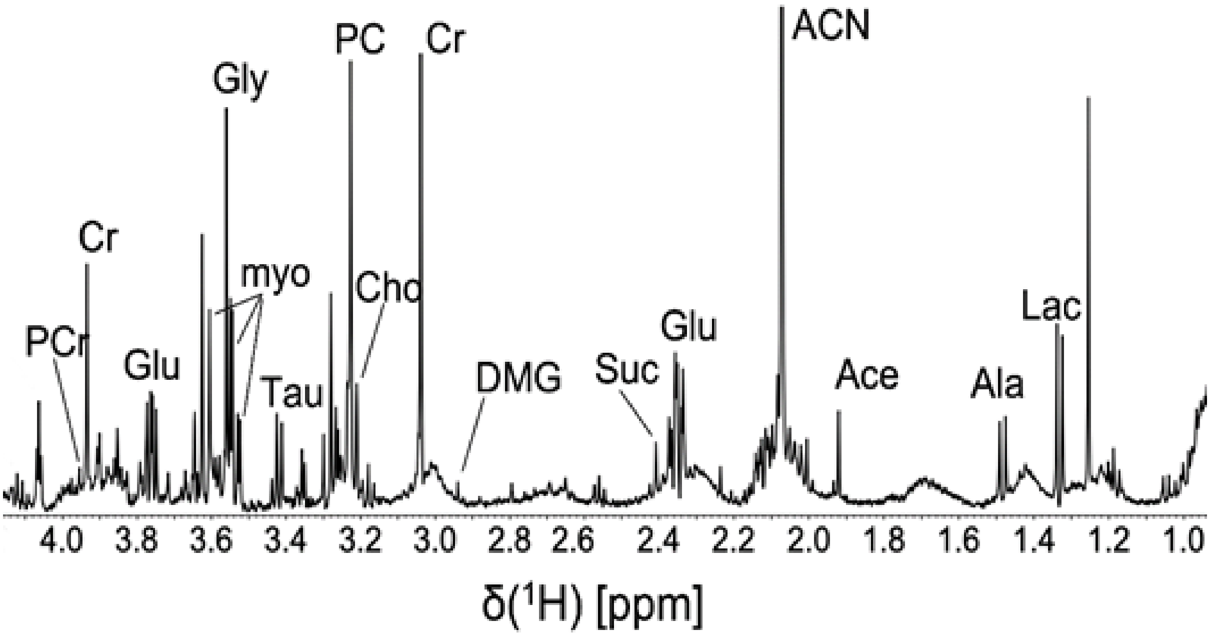


(B)

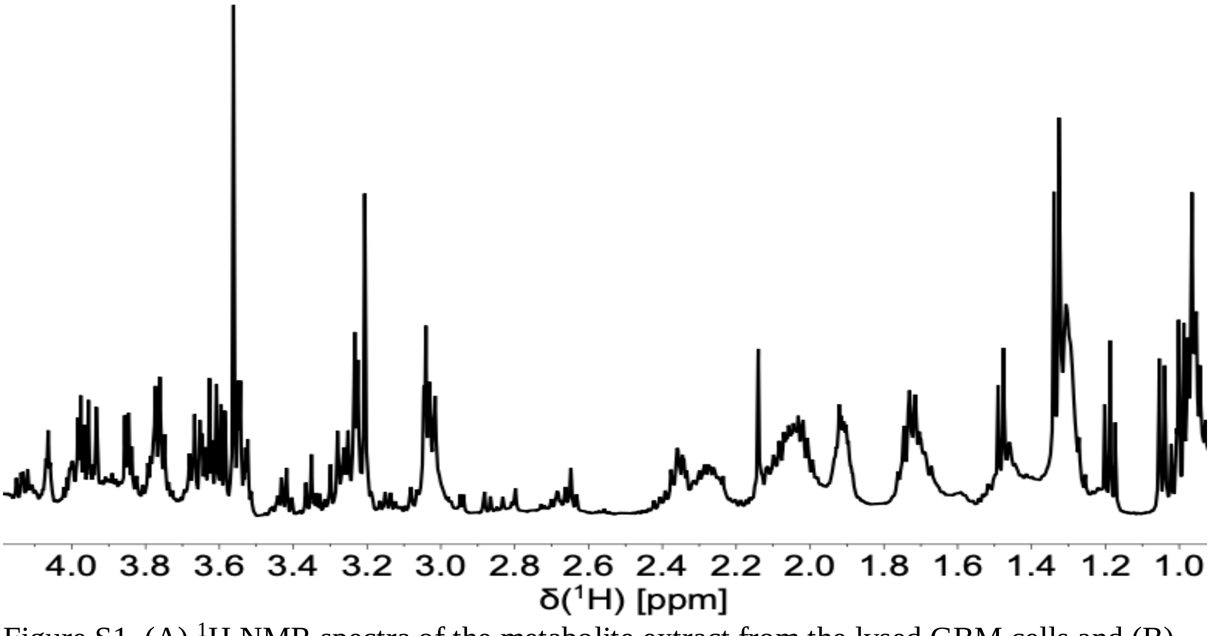


Figure S1. (A) $^{1}$H NMR spectra of the metabolite extract from the lysed GBM cells and (B) of the whole, intact cells sample.

(A)

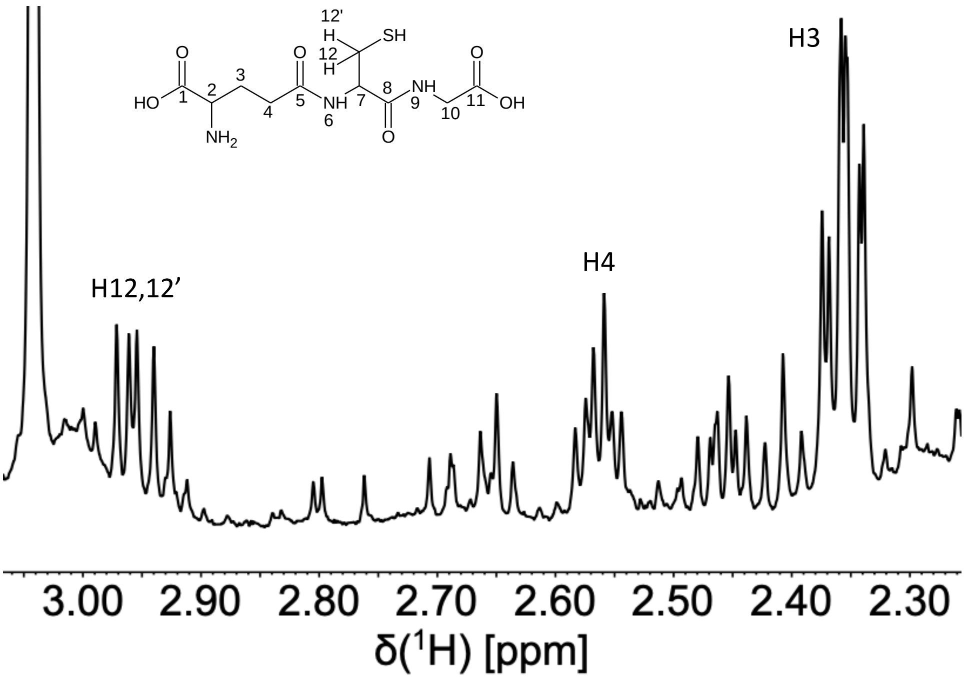


(B)

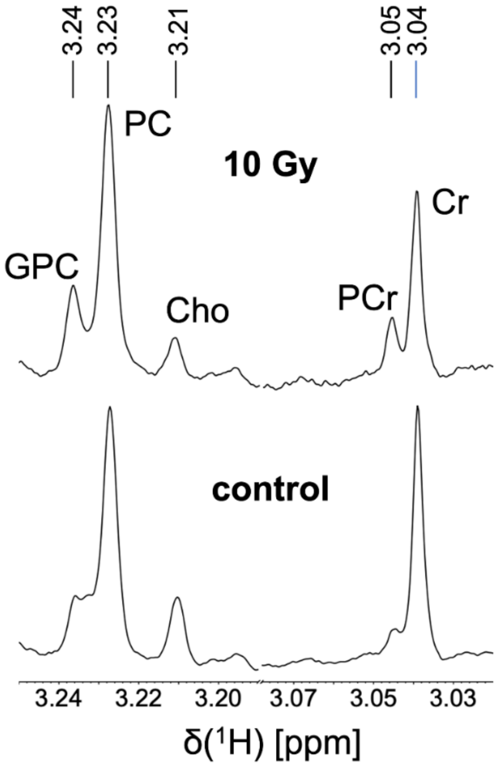


Figure S2. (A) $^{1}$H NMR spectrum of the metabolite extract from a lysed sample consisting of a very large number of cells (>40 M) where glutathione (GSH) was detected and (B) Spectra detail to indicate the level of spectral resolution; a comparison between non-irradiated cells and 10 Gy-irradiated cells is provided.

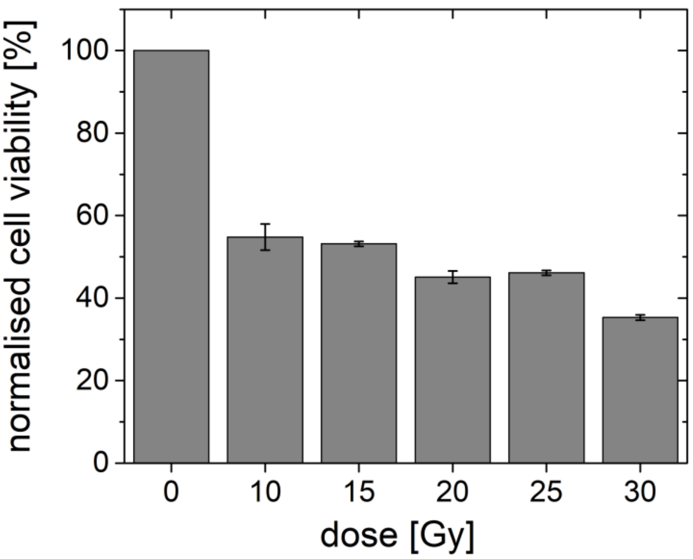


Figure S3. Cell viability as a function of the dose, measured by the metabolic activity quantification (MTT test).

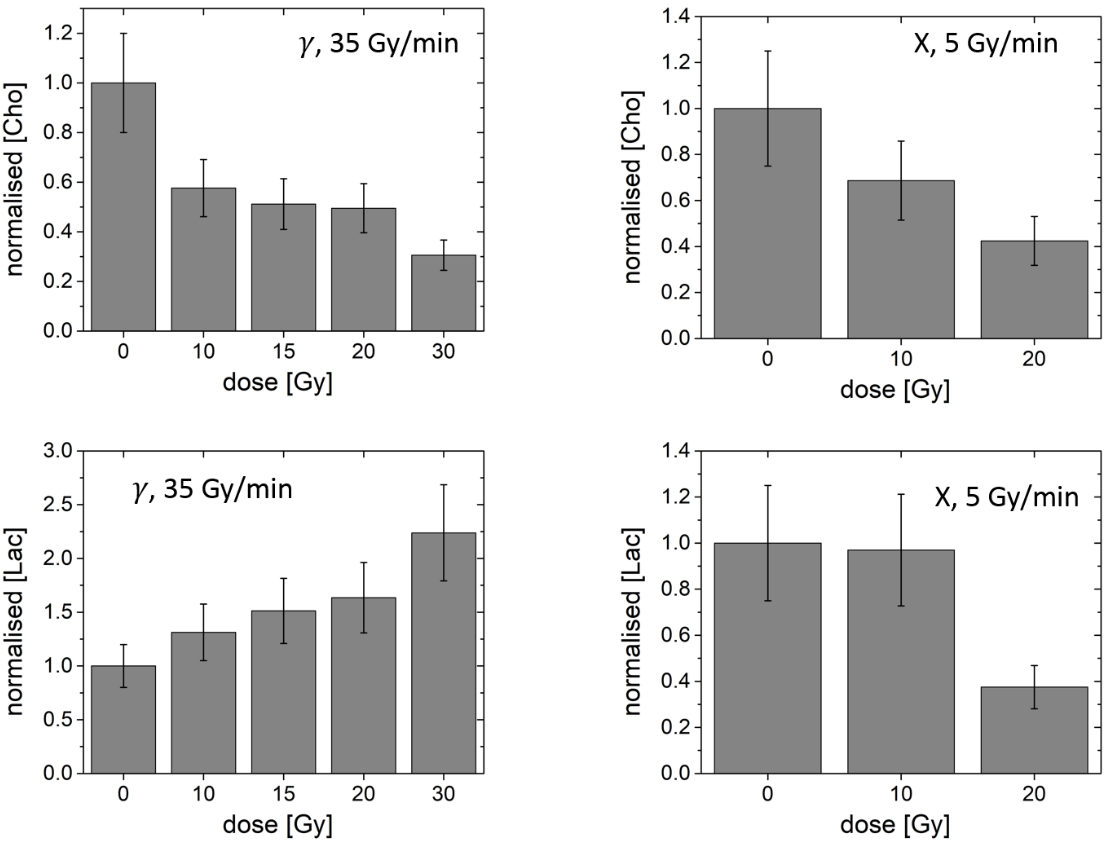


Figure S4. [Cho] and [Lac] as a function of radiation dose, normalized to the sum of signal integrals of the detected metabolites in order to verify the robustness of the normalization.

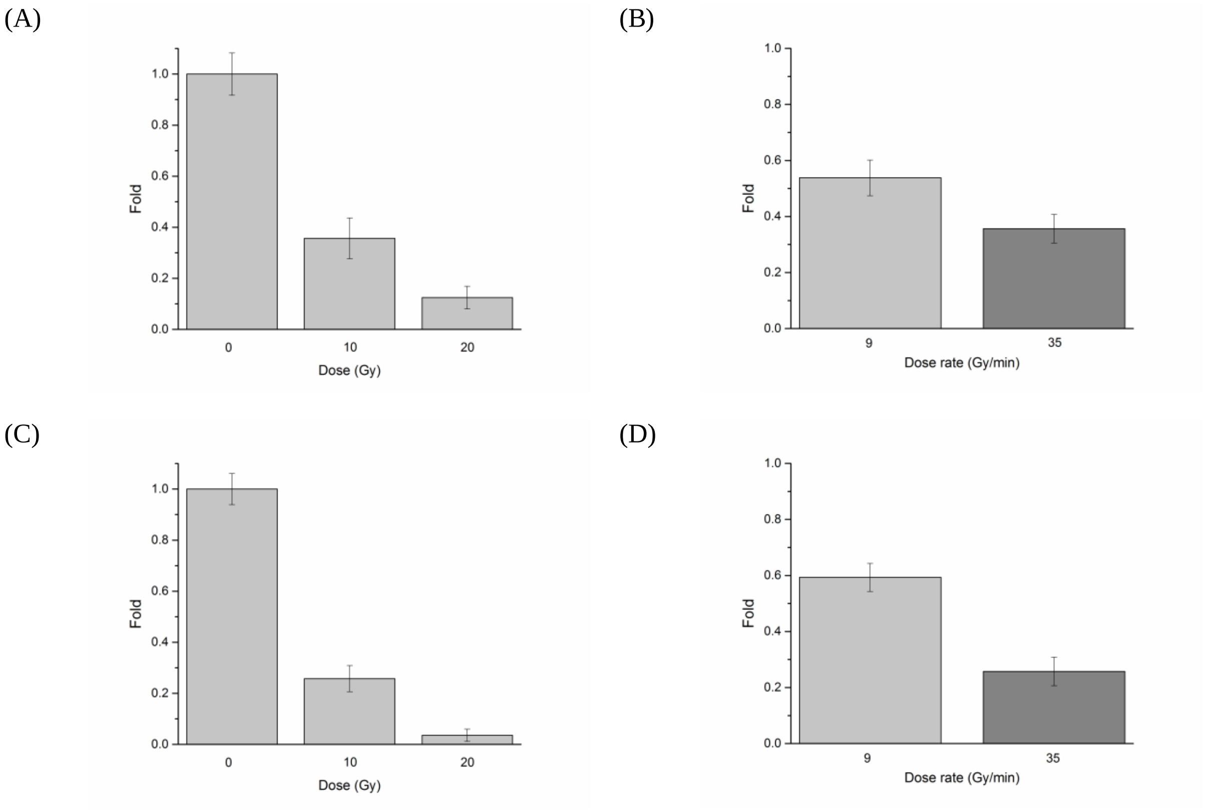


Figure S5. Effects of the irradiation dose (A, C) and dose-rate (B, D) on the glioblastoma cells, assessed within two hours after irradiation by HPLC: sarcosine (A, B) and cystine (C, D) are detected biomarkers. The concentrations of each analyte were normalized to the first point (unirradiated). Chromatograms were obtained using a Reversed Phase Liquid Chromatography system with o-phthaldehyde and 9-fluorenylmethoxycarbonyl chloride for online pre-column derivatization and UV-Vis detection at 241 and 330 nm.

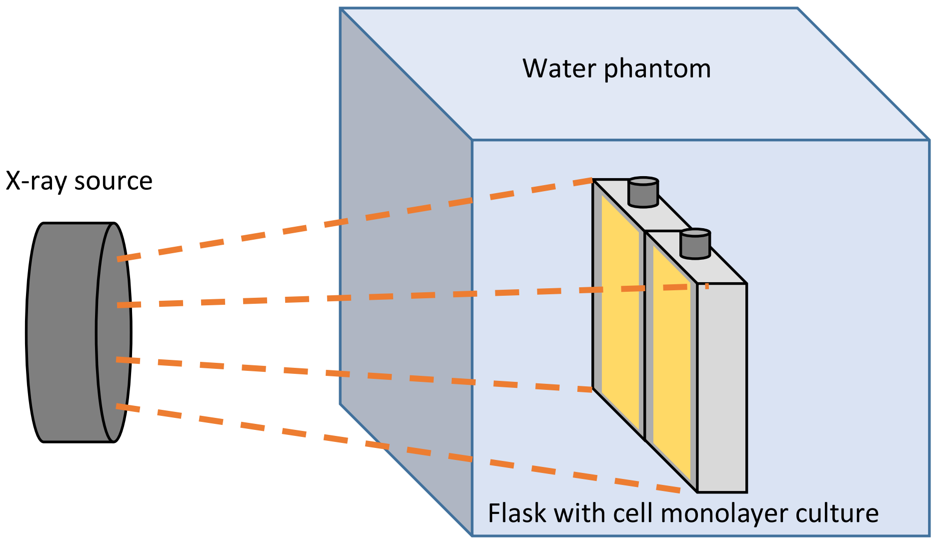


Figure S6. Scheme of the experimental set-up for irradiation of cell samples in the clinical X-ray irradiator. The distance between the X-ray source and the flasks (each 0.1x0.07x0.04 m) was d = 1.2 m and a water phantom cube with the side of 0.4 m was used for flask fixation.
